# An *rpw8* quadruple mutant of Arabidopsis Col-0 is partially compromised in bacterial and fungal resistance

**DOI:** 10.1101/839308

**Authors:** Baptiste Castel, Yan Wu, Shunyuan Xiao, Jonathan D G Jones

**Affiliations:** The Sainsbury Laboratory, University of East Anglia, Norwich Research Park, NR4 7UH Norwich, United Kingdom; Institute for Bioscience and Biotechnology Research & Department of Plant Sciences and Landscape Architecture, University of Maryland College Park, Rockville, MD 20850, USA

## Abstract

The plant immune system relies on both cell-surface and intracellular NLR (nucleotide-binding, leucine-rich repeat) receptors. NLRs respond to pathogen effectors and activate effector-triggered immunity: a cocktail of responses, often accompanied by cell death, resulting in resistance.

*RPW8* encodes an unusual non-NLR Resistance (R) protein and confers broad-spectrum powdery mildew resistance. It requires genetic components also required by some NLRs, resembles the HeLo-containing protein MLKL (necroptosis executor in animals) and HET-S (cell death executor in fungi) and is targeted to the extra-haustorial membrane during powdery mildew infection by its N-terminal non-cleaved signal anchor domain. RPW8 displays extensive recent duplication events in Arabidopsis and certain alleles can induce oligomerisation-dependent activation of the NLR RPP7.

All these features enabled us to formulate hypotheses for RPW8 function: (1) RPW8 could be a cell death executor for defence against pathogens. (2) RPW8 could be a decoy for effector targets.

To test these hypotheses, we generated a quadruple knock-out mutant of the four *RPW8*-homologous copies in Arabidopsis Col-0, using CRISPR. The mutant still displays cell death upon activation of four well-characterised NLRs. However, it is partially impaired in powdery mildew resistance and also in bacterial resistance. Interestingly Col-0_*rpw8* is delayed in flowering transition. In conclusion, RPW8 plays a broad role in immunity and plant development, beyond resistance to powdery mildew. There is no evidence that it is involved in executing ETI-associated cell death.

## Introduction

Plants have co-evolved with their pathogens for millions of years, driving natural selection for effective immunity. The plant immune system relies on recognition of conserved pathogen-, microbe- or damage-associated molecular patterns (PAMPs, MAMPs and DAMPs) by pattern recognition receptors (PRRs) generally localised at the cell surface. PRR activation leads to Ca^2+^ influx, mitogen activated protein kinase (MAPK) activation, reactive oxygen species (ROS) production and transcriptional reprogramming, resulting in PAMP-triggered immunity (PTI) (Couto & Zipfel, 2016). Pathogens evolved effectors to colonize plants. Some effectors interfere with PTI, resulting in effector-triggered susceptibility (ETS). In turn, plants evolved *Resistance*- (*R*-) genes that recognise effectors. *R*-gene products (*i.e.* R-proteins) are mainly intracellular. Their activation results in effector-triggered immunity (ETI) which is generally accompanied by salicylic acid production and the hypersensitive response (HR, not to be confused with *HR1*, *HR2*, *HR3* and *HR4*, which are “*Homologues of RPW8*” Arabidopsis genes), a form of programmed cell death at the site of infection (Jones & Dangl, 2006). Elevation of salicylic acid levels induces local defence and systemic acquired resistance (Durner *et al.*, 1997). Localized HR is thought to stop propagation of the pathogen within the host (Morel & Dangl, 1997).

Investigation of the plant immune system revealed hundreds of *R*-genes in most angiosperm genomes. Most of them belong to the Nucleotide-Binding (NB), Apaf-1, R-protein and CED-4 (ARC) and Leucine-rich repeat (LRR) (NLR) family (Kourelis & Van Der Hoorn, 2018). Earlier this year, cryo-electron microscopy was employed to resolve the structure of ZAR1, a plant NLR, in its non-active and active form (Dangl & Jones, 2019). The non-active form binds ADP in its NB domain and is monomeric (Wang *et al.*, 2019b). The active form binds ATP in its NB domain and is pentameric, forming a resistosome (Wang *et al.*, 2019a) that imposes induced proximity on its N-terminal region. The function of the resistosome is to activate immunity, but the mode of action is not known.

Some components genetically required for NLR function have been characterised. For instance, EDS1 (Enhanced Disease Susceptibility 1) and PAD4 (Phytoalexin Dependent 4) are lipase-like proteins, conserved between many plants and required for TIR-NLR-mediated immunity (Wiermer *et al.*, 2005). TIR-NLRs form a sub-family of plant NLRs that is defined by the presence of an N-terminal TIR (Toll-like, Interleukin-1 receptor and R-protein) domain and specific motifs in the NB-ARC domain. EDS1 is also required redundantly with ICS1 (Isochorismate Synthase 1, required for salicylic acid biosynthesis) for some CC-NLRs (Venugopal *et al.*, 2009). CC-NLRs form a second sub-family of plant NLRs. They are defined by the presence of an N-terminal CC (coiled-coil) domain and specific motifs in the NB-ARC domain.

*RPW8* (*Resistance to Powdery Mildew 8*) is a genetic locus identified in Arabidopsis that confers broad-spectrum resistance to powdery mildew fungi in accession Ms-0. Two homologous *R*-genes, *RPW8.1* and *RPW8.2*, from the same locus contribute to resistance (Xiao *et al.*, 2001). Transgene analysis showed that adequate expression of *RPW8.1* and *RPW8.2* also confers resistance to a virulent strain of the oomycete *Hyaloperonospora arabidopsidis* (*Hpa*), the cause of downy mildew disease (Wang *et al.*, 2007). Transgenic tobacco plants expressing *RPW8.1* and *RPW8.2* showed enhanced resistance to powdery mildew (Xiao *et al.*, 2007). More recently, heterologous expression of *RPW8.1* in rice and *RPW8.2* in grapevine has also been shown to increase resistance to *Magnaporthe oryzae* (Li *et al.*, 2018) and powdery mildew (Hu *et al.*, 2018), respectively. Both *RPW8.1* and *RPW8.2* (noted as *RPW8* in later text, unless indicated otherwise) encode a small protein (~20 kDa) with a predicted CC domain. Intriguingly, RPW8 shares sequence homology to the N-termini of a third sub-family of plant NLRs, called RPW8-NLRs. This sub-family of NLRs often have only a few members but exists in almost all plants (Zhong & Cheng, 2016). At least in angiosperms, these RPW8-NLRs are helper NLRs required for signalling of multiple sensor NLRs (Peart *et al.*, 2005; Bonardi *et al.*, 2011; Dong *et al.*, 2016; Qi *et al.*, 2018; Adachi *et al.*, 2019; Castel *et al.*, 2019a; Wu *et al.*, 2019).

Genetic analysis showed that the resistance function of RPW8 requires previously characterised immune components, including EDS1, PAD4 and the salicylic acid pathway, but does not require NDR1 (which is usually required for CC-NLRs), nor COI1 and EIN2 (which regulate jasmonic acid and ethylene signalling respectively) (Xiao *et al.*, 2005). Because EDS1, PAD4 and salicylic acid are generally associated with TIR-NLR-mediated immunity, the above results imply a functional link between RPW8 and TIR-NLR signalling.

During their co-evolution with hosts, some filamentous pathogens, such as powdery mildew and rust fungi, and oomycetes have evolved a common invasive strategy: the formation of haustoria as their feeding structures. Recent studies have shown that the haustorium is encased by a host-derived special membrane called the extra-haustorial membrane (EHM). Interestingly, RPW8.2 is specifically targeted to the EHM during powdery mildew infection (Wang *et al.*, 2009, 2013). The N-terminal domain of RPW8.2, along with two EHM-targeting motifs, is predicted to associate with membranes, and is required for EHM targeting and resistance function of RPW8.2. Intriguingly, the N-terminal portion (~90 amino acids) of RPW8 shares similarity with MLKL from animals and HELLP and other HeLo-domain-containing fungal proteins (Daskalov *et al.*, 2016). MLKL can oligomerise to form pores at the membrane during necroptosis in animals (Murphy *et al.*, 2013). HELLP has a similar function in fungi and can form prions via its C-terminus. Prionisation is associated with membrane targeting and disruption, followed by cell death. A glycine zipper motif conserved between MLKL, HELLP and RPW8 is required for HELLP membrane targeting (Daskalov *et al.*, 2016). In parallel, RPW8 is capable of inducing HR (Xiao *et al.*, 2001) and overexpression of RPW8 can lead to cell death (Xiao *et al.*, 2003, 2005). However, it is not known whether RPW8’s membrane-targeting and cell death induction share the same molecular basis with that of HELLP and MLKL.

Another interesting feature of the *RPW8* locus is the intraspecific gene amplification and diversity within Arabidopsis. RenSeq was conducted on 65 Arabidopsis accessions and revealed two types of sequence variation at the *RPW8* locus (Barragan *et al.*, 2019; Van de Weyer *et al.*, 2019). Firstly, there have been a large number of recent duplication events. There are seven sub-clades of *RPW8* paralogs: *RPW8.1*, *RPW8.2*, *RPW8.3*, *HR1*, *HR2*, *HR3* and *HR4*. Each Arabidopsis accession contains a unique combination of these alleles with different copy numbers. For instance, Ms-0 contains five *RPW8* paralogs: *RPW8.1* and *RPW8.2*, which confer resistance to powdery mildew, and *HR1*, *HR2* and *HR3*, which are not required for powdery mildew resistance (Xiao *et al.*, 2001). Col-0, which is susceptible to powdery mildew, contains *HR1*, *HR2*, *HR3*, and *HR4*, an additional homolog most closely related to RPW8.1 (Xiao *et al.*, 2001, 2004; Berkey *et al.*, 2017). Interestingly, TueWa1-2 contains as many as 13 *RPW8* paralogs (including three copies of *RPW8.1*, two of *RPW8.2*, four of *RPW8.3*, one of *HR1*, one of *HR2*, one of *HR3* and one of *HR4*). The selective advantage of such extreme variation in duplication events is not understood. Secondly, some *RPW8* genes encode proteins with intragenic C-terminal duplications. For instance, RPW8.1 from some accessions contains one or more duplications of a 21-amino acid motif (Orgil *et al.*, 2007), and HR4 from Col-0 contains five duplications of a 14-amino acid motif (Xiao *et al.*, 2004), whereas HR4 from ICE106 contains only two, and HR4 from Fei-0 only one such motif (Barragan *et al.*, 2019).

A large-scale genetic analysis on intraspecific hybrid necrosis revealed that some *RPW8* alleles are “incompatible” with alleles of an NLR-type *R*-gene at the *RPP7*-locus (Chae *et al.*, 2014). Some Arabidopsis crosses display spontaneous necrosis in F1, due to the presence of two incompatible alleles, one coming from each parent. The “*Dangerous Mix*” (*DM*) loci have been mapped. Particularly, *DM6* (*RPP7*) and *DM7* (*RPW8*) are responsible for incompatibility in three crosses: Mrk-0 × KZ10, Lerik1-3 × Fei-0 and ICE79 × Don-0. In KZ10 and Fei-0, the *RPW8* genes responsible for the phenotype are *RPW8.1* and *HR4* respectively (Barragan *et al.*, 2019). RPW8.1^KZ10^ has three C-terminal intragenic repeats, the maximum observed among known RPW8.1 alleles. Interestingly, Arabidopsis accessions carrying an RPW8.1 allele with three C-terminal repeats all form a dangerous mix with Mrk-0. In contrast, HR4^Fei-0^ has only one C-terminal repeat. Arabidopsis accessions carrying an HR4 allele with one C-terminal repeat all form a dangerous mix with Lerik1-3. The number of C-terminal repeats thus correlates with incompatibility with different RPP7 alleles. However, the function of the repeat is yet to be discovered. In addition, HR4^Fei-0^ induces oligomerisation of RPP7b^Lerik1-3^, resulting in HR in *Nicotiana benthamiana* (Li *et al.*, 2019).

We can formulate two hypotheses about RPW8 function. (1) Based on sequence similarity, RPW8 could function as the HeLo domain-containing proteins MLKL, HET-S and HELLP from animals and fungi. This function would be a ligand-dependent oligomerization resulting in membrane targeting and disruption for programmed cell death. (2) RPW8 could also be a decoy of effector targets. NRG1 and ADR1 are low copy number and are helper NLRs (Jubic *et al.*, 2019). Their conserved RPW8 domain could in theory provide a potent effector target to suppress immunity mediated by multiple sensor NLRs. RPW8 paralogs could encode decoys for such effectors. They could either trap the effectors to prevent their virulence function or be modified and guarded by NLRs such as RPP7. The extreme degree of recent duplication of RPW8 in Arabidopsis is consistent with this hypothesis.

To test both hypotheses, we generated a quadruple *rpw8* mutant (*hr1-hr2-hr3-hr4*, we called Col-0_*rpw8*) in Arabidopsis Col-0 to test for possible altered immunity. Our results indicate a function for RPW8 beyond defence against powdery mildew pathogens.

## Materials and Methods

### Plant genotypes and growth conditions

*Arabidopsis thaliana* (Arabidopsis) accessions used in this study is Columbia-0 (Col-0). Col-0_*eds1* is an *eds1a-eds1b* double CRISPR mutant, published as *eds1-12* (Ordon *et al.*, 2017). Seeds were sown directly on compost and plants were grown at 21°C, with 10 hours of light and 14 hours of dark, 75% humidity. For seed collection, 5-week old plants were transferred under long-day condition: 21°C, with 16 hours of light and 8 hours of dark, 75% humidity. For *Nicotiana benthamiana, s*eeds were sown directly on compost and plants were grown at 21°C, with cycles of 16 hours of light and 8 hours of dark, 55% humidity. *N. benthamiana_nrg1* is an *nrg1* CRISPR mutant of *N. benthamiana* (Castel *et al.*, 2019a).

### Generation of an *hr1-hr2-hr3-hr4* (aka *rpw8*) null mutant in Arabidopsis Col-0 using CRISPR

CRISPR modules were assembled using the Golden Gate cloning method. The detailed Golden Gate method and protocols can be found in (Engler *et al.*, 2009, 2014; Weber *et al.*, 2011). Specific details for CRISPR module assembly can be found in (Castel *et al.*, 2019b). All the vector used can be found on Addgene (addgene.org). 12 sgRNAs targeting *HR1* (TGGCGTCGTGAAGGAGTTGG[nGG], CGACGCCATCATAAGAGCCA[nGG] and GTTCATCGACTTCTTCGGTG[nGG]), *HR2* (TGCTCTCCAAATCCTTCACG[nGG], GTCTCGATTCTACAATCTTG[nGG] and GGTTCTTGTCGAAGCTTATG[nGG]), *HR3* (GGTTAGTGAGATTATGGCAG[nGG], GTCTTGATGCTACAATCTTT[nGG] and CGATAAGCTTAGCGAAGAAG[nGG]) and *HR4* (GCTTGCTGTAATCAAAACAG[nGG], TGGAAAGTATCAGTCCGGTG[nGG] and GGAGACGCGTAAGACTTTCG[nGG]) were designed and assembled by PCR to a sgRNA backbone and 67 bp of the *AtU6-26* terminator. sgRNA1 to sgRNA12 were assembled with the *AtU6-26* promoter in the Golden Gate compatible level 1 *pICH47761*, *pICH47772, pICH47781, pICH47791, pICH47732, pICH47742, pICH47751*, *pICH47761*, *pICH47772, pICH47781, pICH47791* and *pICH47732* respectively. A level 1 vector *pICH47811* (with expression in reverse orientation compared to the other level 1 modules) containing a human codon optimized allele of *Cas9* under the control of the *AtRPS5a* promoter and the *Pisum sativum rbcS E9* terminator (Addgene: 117505) was assembled with a FAST-Red selectable marker (Addgene: 117499) into a level M vector *pAGM8031*, using the end-linker *pICH50892*. sgRNA1 to sgRNA5 were assembled into a level M vector *pAGM8067*, using the end-linker *pICH50872*. sgRNA7 to sgRNA9 were assembled into a level M vector *pAGM8043*, using the end-linker *pICH50914*. sgRNA10 to sgRNA12 were assembled into a level M vector *pAGM8043*, using the dummy module *pICH54033* and the end-linker *pICH50881*. The four level M modules were assembled into level P vector *pICSL4723-P1*, using the end-linker *pICH79264*. Level 1 vectors were cloned using BsaI enzyme and carbenicillin resistance. Level M vectors were cloned using BpiI enzyme and Spectinomycin resistance. Level P vector was cloned using BsaI enzyme and kanamycin resistance. The final vector map can be found in the Supplemental Information. It was expressed via *Agrobacterium tumefaciens* strain GV3101 in Arabidopsis Col-0. In the first generation after transformation, we recovered somatic mutants. The non-transgenic progeny of a somatic mutant contained a homozygous quadruple knock-out line. One mutation is a 6579 bp deletion between C173 of *HR1* and G22 of *HR3*. The second mutation is c.28delA in *HR4*, causing an early stop codon at the N-terminal encoding region. The resulting line lacks functional *HR1*, *HR2*, *HR3*, *HR4* and is T-DNA free. We called this line Col-0_*rpw8*.

### Transcript level measurement

For gene expression analysis, RNA was isolated from three biological replicates and used for subsequent reverse transcription quantitative PCR (RT-qPCR) analysis. RNA was extracted using the RNeasy Plant Mini Kit (QIAgen) and treated with RNase-Free DNase Set (QIAGEN). Reverse transcription was carried out using the SuperScript IV Reverse Transcriptase (ThermoFisher). qPCR was performed using CFX96 Touch^TM^ Real-Time PCR Detection System. Primers for qPCR analysis of *PR1* are ATACACTCTGGTGGGCCTTACG and TACACCTCACTTTGGCACATCC. Primers for qPCR analysis of *EF1α* are CAGGCTGATTGTGCTGTTCTTA and GTTGTATCCGACCTTCTTCAGG. Data were analysed using the double delta Ct method (Livak & Schmittgen, 2001).

### Cell death assay in Arabidopsis

*Pseudomonas fluorescens* engineered with a Type Three Secretion system (Pf0-1 EtHAn) (Thomas *et al.*, 2009) carrying pBS46:AvrRps4, pBS46:AvrRps4^KRVY^, pBS46:AvrRpt2, pVSP61:AvrRpm1 or pVSP61:AvrPphB were grown on selective KB-medium agar plate for 48 hours at 28 °C. Bacteria were harvested from plate, re-suspended in infiltration buffer (10 mM MgCl2, pH 5.6) and concentration was adjusted to OD_600_ = 0.2 (~10^8^ cfu/ml). The abaxial surface of 4-week old Arabidopsis leaves were hand-infiltrated with 1 ml needle-less syringe. Cell death was monitored 24 hours after infiltration.

### Cell death assay in *N. benthamiana*

*A. tumefaciens* strains were streaked on selective media and incubated at 28 °C for 24 hours. A single colony was transferred to liquid LB medium with appropriate antibiotic and incubated at 28 °C for 24 hours in a shaking incubator (200 rotations per minute). The resulting culture was centrifuged at 3000 rotations per minute for 5 minutes and resuspended in infiltration buffer (10 mM MgCl2, 10 mM MES, pH 5.6) at OD_600_ = 0.4 (2×10^8^ cfu/ml). For co-expression, each bacterial suspension was adjusted to OD_600_ = 0.4 in the final mix. The abaxial surface of 4-weeks old *N. benthamiana* were infiltrated with 1 ml needle-less syringe. Cell death was monitored three days after infiltration.

Vector used are: RRS1-^RK1221Q^-FLAG and RPS4-HF (Sarris *et al.*, 2015), EDS1-V5 and SAG101-Myc (Huh *et al.*, 2017), NRG1B-HF (Castel *et al.*, 2019a) and HR2-mNeon (this article). *HR2* was amplified from Col-0 (Fw: GGCTTAAUATGCCTCTTACCGAGATTATCG, Rv: AACCCGAUTTCAAAACGAAGCGAATTC) and mNeon c-terminal tag was amplified from pICSL50015 (Addgene: 50318) (Fw: ATCGGGTUTGGTGAGCAAGGGAGAGGAGG, Rv: GGTTTAAUTTACTTGTAAAGCTCGTCCA) using “KAPA HiFi HotStart Uracil+ ReadyMix (2X)” enzyme (KAPABIOSYSTEMS), following the manufacturer protocol. Amplicons were assembled in a USER compatible vector LBJJ234-OD (containing a FAST-Red selectable marker and a 35S / Ocs expression cassette, pre-linearized with PacI and *Nt. BbvcI* restriction enzymes), with the USER enzyme (NEB), following the manufacturer protocol. The final vector map can be found in the Supplemental Information.

### Bacterial growth measurement

*Pseudomonas syringae* pv. tomato strain DC3000 carrying pVSP61:AvrRps4-HA or pVSP61 empty vector were grown on selective KB-medium agar plate for 48 hours at 28 °C. Bacteria were harvested from plate, re-suspended in infiltration buffer (10 mM MgCl_2_, pH 5.6) and concentration was adjusted to OD_600_ = 0.0005 (~2.5×10^5^ cfu/ml). The abaxial surface of 5-week old Arabidopsis leaves were hand-infiltrated with 1 ml needle-less syringe. Plants were covered with a lid for the first 12 hours of bacterial growth. For quantification, leaf samples were harvested with a 6 mm diameter cork-borer, resulting in a ~0.283 cm^2^-sized leaf disc. Two leaf discs per leaf were harvested and used as single sample. For each condition, four samples were collected just after infiltration and six samples were collected 72 hours after infiltration. Samples were ground in 200 μl of infiltration buffer, serially diluted (5, 50, 500, 5000 and 50000 times) and spotted (5 to 10 μl per spot) on selective KB-medium agar plate to grow 48 hours at 28 °C. The number of colonies (cfu per drop) was monitored and the bacterial growth was expressed in cfu/cm^2^ of leaf tissue.

### *Albugo candida* propagation

For propagation of *Albugo candida* race 2V (Rimmer *et al.*, 2000), zoospores were suspended in water (~10^5^ spores/ml) and incubated on ice for 30 min. The spore suspension was then sprayed on plants using a Humbrol® spray gun (~700 μl/plant) and plants were incubated at 4 °C in the dark overnight. Infected plants were kept under 10 hours light (20 °C) and 14 hours dark (16 °C) cycles. Phenotypes were monitored 12 days after spraying.

### Powdery mildew propagation

For testing with powdery mildew, plants were grown under short-day (8 hr light, 16 hr dark) and 75% relative humidity at 22 °C for 7 weeks before inoculation with an adapted powdery mildew isolate *Golovinomyces cichoracearum* (*Gc*) UCSC1 or a non-adapted isolate *Gc* UMSG1 as previously reported (Xiao *et al.*, 2005; Wen *et al.*, 2011). Disease phenotypes with *Gc* UCSC1 were visually scored and photographed. Disease susceptibility was further assessed by counting total number of spores per mg leaf tissue. At least 6 infected leaves were weighed and combined as one leaf sample, and 4 samples were collected from 12 infected plants for disease quantification. A spore suspension of each sample was made by vortexing the leaves for 1 min in 10 ml of H_2_O containing 0.02 % Silwet L-77 and used for spore counting using LunaTM Automated Cell Counter (Logos biosysems). Spore counts were normalized to the fresh weight of the leaf samples. For infection tests with *Gc* UMSG1, which can largely penetrate the cell wall of Arabidopsis but fails to establish micro-colony capable of sporulation (Wen *et al.*, 2011), inoculated leaves were collected at 5 dpi and subjected to trypan blue staining, and total the hyphal length was measured as previously reported (Wen *et al.*, 2011). All infection trials were repeated three times with similar results.

### Phylogenetic reconstruction of RPW8 homologs in Arabidopsis Col-0

RPW8 domain boundaries were predicted using SMART (http://smart.embl-heidelberg.de/) (Letunic & Bork, 2017). Amino acid sequences of RPW8 domains were aligned using the MUSCLE method. The evolutionary history was inferred using the Neighbor-Joining method. The evolutionary distances were computed using the Poisson correction method. All positions with less than 95% site coverage were eliminated. Evolutionary analyses were conducted using the software MEGA7.

## Results

### CRISPR mutagenesis enables recovery of an *rpw8* quadruple mutant

In Col-0, the four *RPW8* homologs *HR1*, *HR2*, *HR3* and *HR4* are located in an 11.8 kb cluster on Chromosome 3 (**Figure 1**). To generate null alleles, we designed three sgRNAs per gene, targeting their N-terminal encoding region. Cas9 activity at the sgRNA target could result in large deletion of several genes and/or indels causing early stop codons in individual genes. We expressed the 12 sgRNAs along with Cas9 from a single T-DNA in Arabidopsis Col-0. We identified a line lacking functional *HR1*, *HR2*, *HR3*, *HR4* that is T-DNA free (see Materials and Methods for more details). We called this line Col-0_*rpw8*.

**Figure 1:**
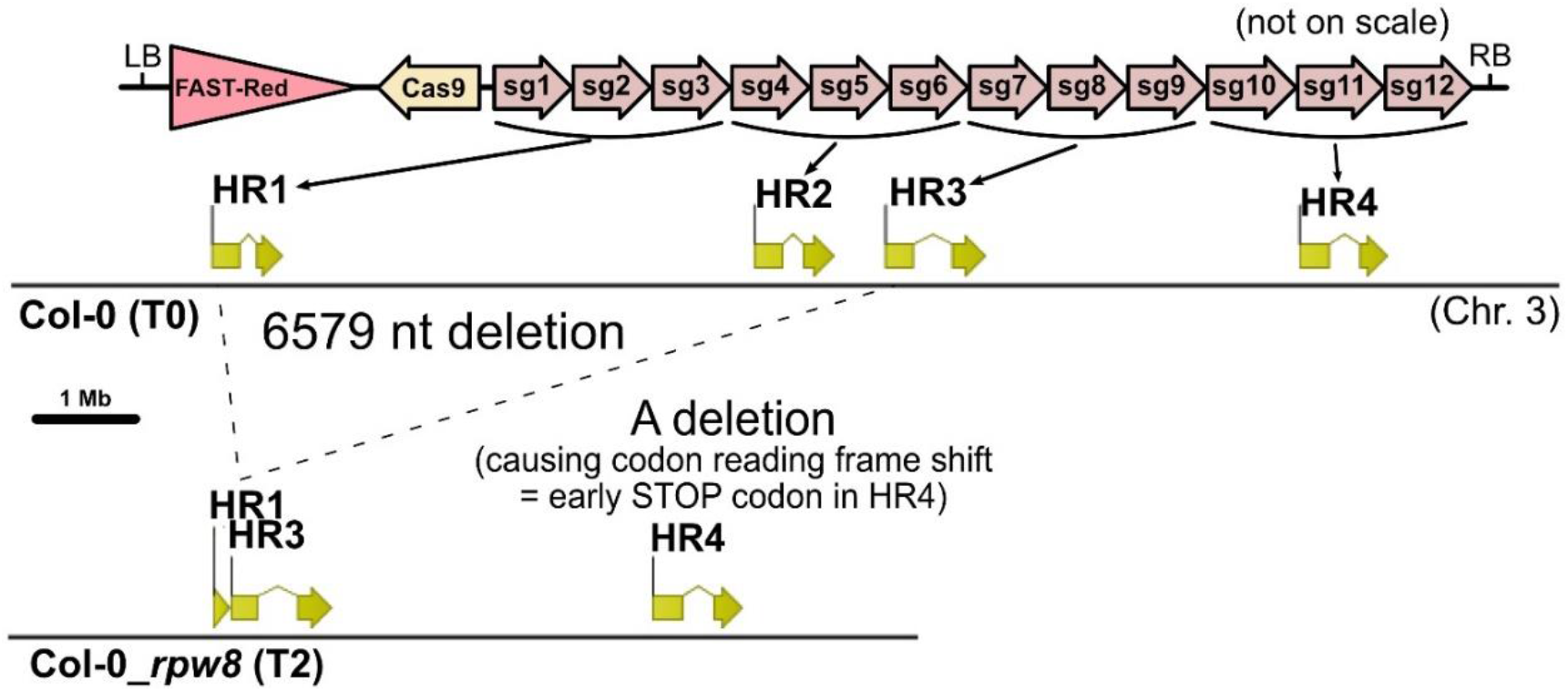
Generating an *hr1-hr2-hr3-hr4* (*rpw8*) null mutant in Arabidopsis Col-0 using CRISPR. *Cas9* (expressed using the *RPS5a* promoter and *E9* terminator) was assembled with 12 sgRNAs (under the *U6-26* promoter and terminators) and the FAST-Red selectable marker using the Golden Gate cloning method. Expression of the construct in Col-0 WT resulted in a 6579 bp deletion between C173 of *HR1* and G22 of *HR3* and a c.28delA in *HR4*, causing an early stop codon at the N-terminal encoding region. LB, RB: T-DNA left and right borders. sg: sgRNA. T0: wild type Col-0. T2: Second generation after transformation. Chr. 3: chromosome 3. Construct map is not on scale. *RPW8* locus cartoons are on scale.

### *rpw8* is slightly autoimmune

Under normal growth conditions, Col-0_*rpw8* plants appear slightly smaller than WT plants (**Figure 2A**). Mutations that result in activation of constitutive defence often display a dwarf phenotype (van Wersch *et al.*, 2016). The “autoimmunity” is usually associated with elevated salicylic acid (SA) levels, resulting in elevated expression of the SA marker gene *PR1*. We measured *PR1* expression in Col-0_*rpw8* to test for autoimmunity. *PR1* is significantly more highly expressed in three independent Col-0_*rpw8* lines than in WT (**Figure 2B**). However, it is only expressed at ~0.6% of the level of *EF1α*. In contrast, activation of the NLR pair RRS1/RPS4 induces *PR1* to ~12× the level of *EF1α* (Castel *et al.*, 2019a). Autoimmune Arabidopsis mutants expressing the effector hopZ5 show *PR1* expression levels of ~3 to ~260 time more than *EF1α* (Jayaraman *et al.*, 2017). Thus, the autoimmunity of Col-0_*rpw8* is detectable but very low.

**Figure 2:**
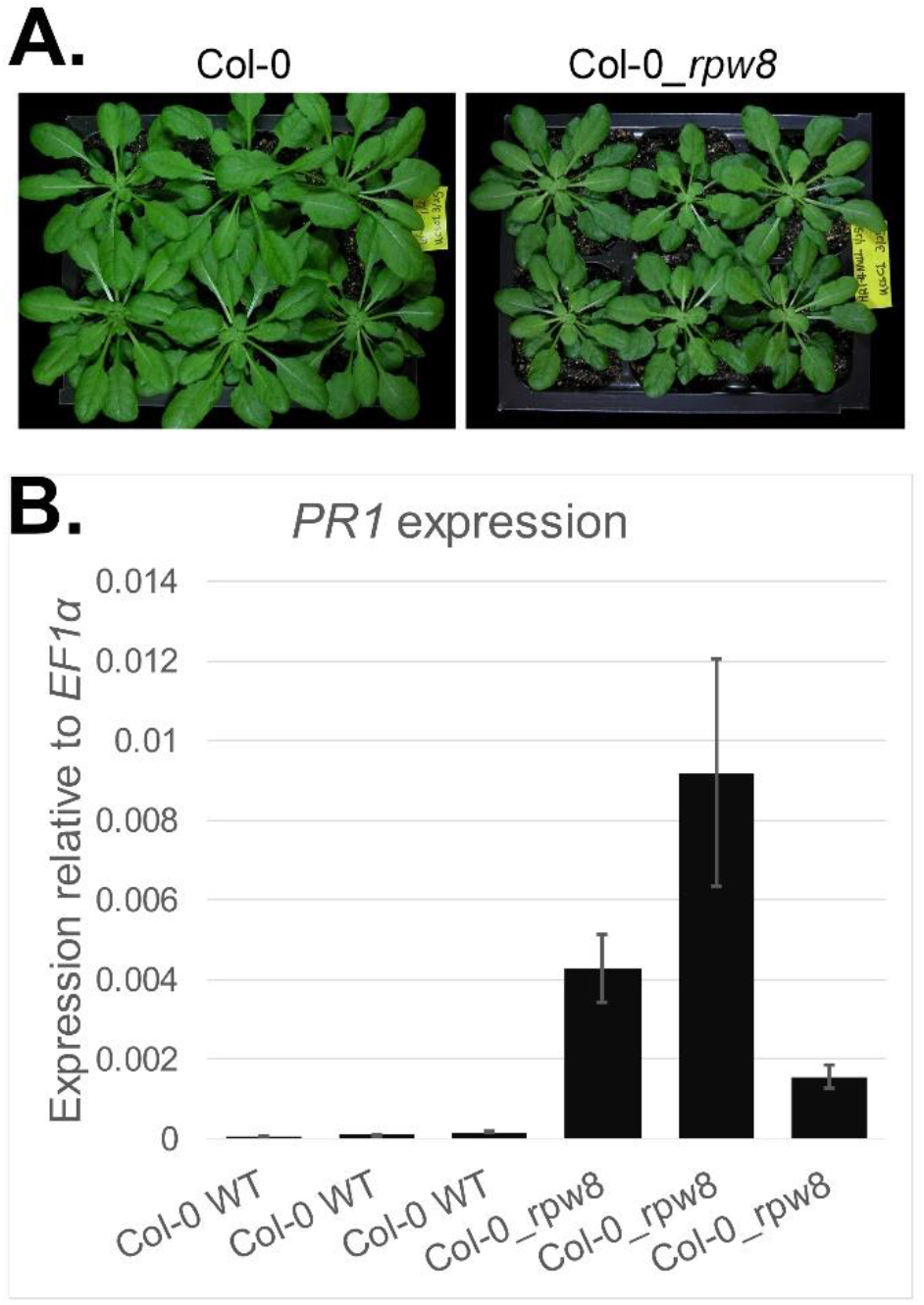
Autoimmunity of Col-0_*rpw8* is detectable but very low. **A.** Col-0_*rpw8* plants are slightly smaller than Col-0 WT. Pictures were taken seven weeks after germination. **B.** *PR1* expression is elevated in three independent Col-0_*rpw8* lines. The expression level is low compared to the *PR1* expression levels in a strong autoimmune mutant or upon NLR activation of WT plants (Jayaraman *et al.*, 2017; Castel *et al.*, 2019a). Each bar represents an individual plant. Error bars represent standard error from three technical replicates. Each of three individual Col-0_*rpw8* replicate is significantly different than each of three Col-0 WT replicate (paired two-tailed student test, p-value < 0.05).

Intriguingly, we found that after growing in short-day (8 hr light, 16 hr dark) conditions for 13 weeks, Col-0 plants started bolting, but there was no sign of bolting in Col-0_*rpw8* plants (**Figure 3**). However, there was no noticeable difference among plants of these two genotypes when they were grown in short-day for 4-6 weeks and then shifted to long-day (16 hr light and 8 hr dark). This unexpected result implies that RPW8 homologs, or perhaps salicylic acid levels, play a role in promoting flowering under short-day conditions.

**Figure 3:**
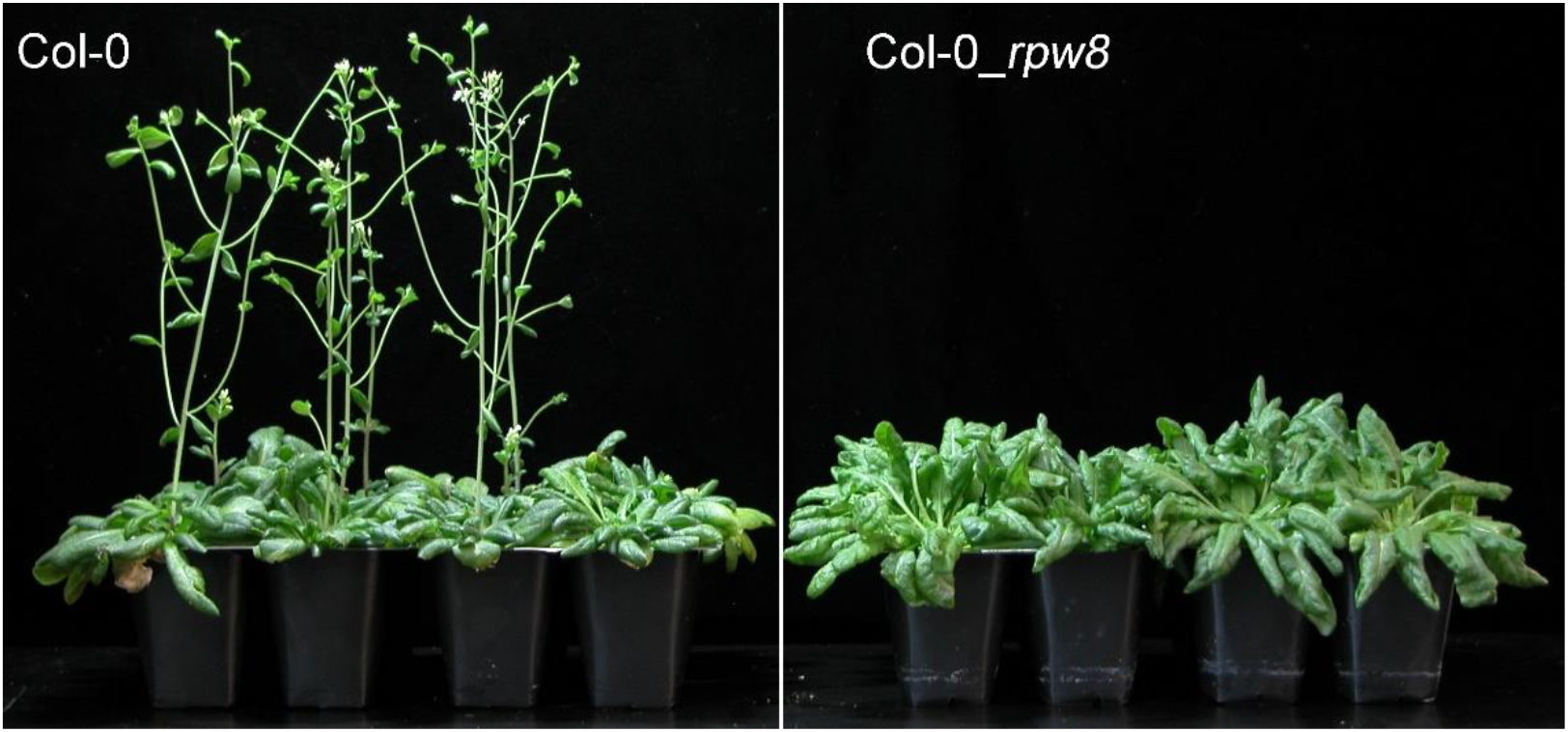
Col-0_*rpw8* exhibits later flowering under short-day conditions. Plants of Col-0 and Col-0_*rpw8* were grown under short-day (8 hr light, 16 hr dark) for 13 weeks. While most (~75%) of Col-0 plants bolted and the remaining initiated bolting, Col-0_*rpw8* plants showed no sign of bolting. Photos were taken at 95 days after seed germination.

### RPW8 homologs are not required for cell death mediated by four well-described NLRs

NLR activation often results in a form of cell death called the hypersensitive response (HR). The bacterial effectors AvrRpm1, AvrRpt2, AvrPphB and AvrRps4 can cause an RPM1-, RPS2-, RPS5- and RRS1/RPS4-dependent HR (Kunkel *et al.*, 1993; Grant *et al.*, 1995; Warren *et al.*, 1998; Gassmann *et al.*, 1999) respectively. We delivered these effectors into Col-0 WT and Col-0_*rpw8*, using Pf0-1, a non-pathogenic strain of *P. fluorescens* engineered with a Type III Secretion System to deliver an effector of interest into the host cell cytosol (Thomas *et al.*, 2009). AvrRps4^KRVY^, an inactive form of AvrRps4, was used as a negative control. Each effector (apart from AvrRps4^KRVY^) can still trigger HR in Col-0_*rpw8* (**Figure 4**). These data indicate that RPW8 homologs are not required for RPM1-, RPS2-, RPS5- or RRS1/RPS4-mediated HR.

**Figure 4:**
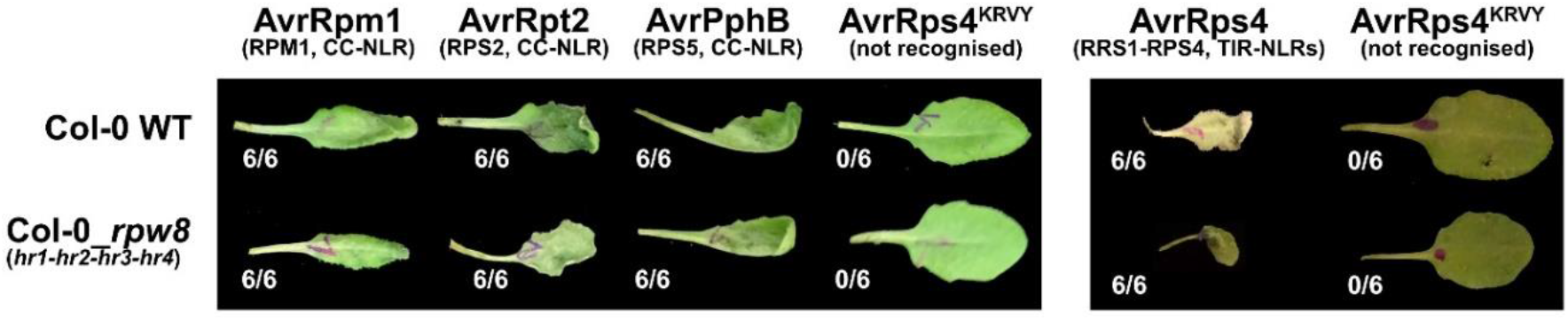
RPW8 homologs are not required for RPM1-, RPS2-, RPS5- or RRS1/RPS4-mediated HR. Effectors were delivered using the Pf0-1 system into leaves of 4-week old plants, OD_600_ = 0.2, pictures taken 24 hpi. Six plants were tested for each combination. Numbers indicate the number of leaves displaying HR. The cognate NLRs (CC- or TIR-) are indicated in parentheses. AvrRps4^KRVY^ is a non-recognised allele of AvrRps4, used as negative control. AvrRps4 was tested in a different day as the other effectors, hence represented in a separate panel.

We then tested whether RPW8 is sufficient to transduce RRS1/RPS4 signal for HR. EDS1, SAG101 and NRG1 are the three major components of TIR-NLR signalling for HR in Arabidopsis and in *Nicotiana benthamiana* (Lapin *et al.*, 2019). We transiently expressed RRS1-R^K1221Q^ (an RPS4-dependent auto-active form of RRS1-R) and RPS4 with EDS1 and/or SAG101 and/or NRG1B and/or HR2 from Arabidopsis. The minimal requirement to reconstruct RRS1/RPS4-mediated HR in *N. benthamiana* is EDS1/SAG101/NRG1, but HR2 is dispensable (**Figure 5 and** **S1**). This indicates that Col-0 RPW8 is neither necessary nor sufficient to transduce RRS1/RPS4 signal for HR. Parenthetically, it also indicates that *N. benthamiana* alleles of SAG101/EDS1/NRG1 cannot transduce the RRS1/RPS4 response, while Arabidopsis alleles can.

**Figure 5:**
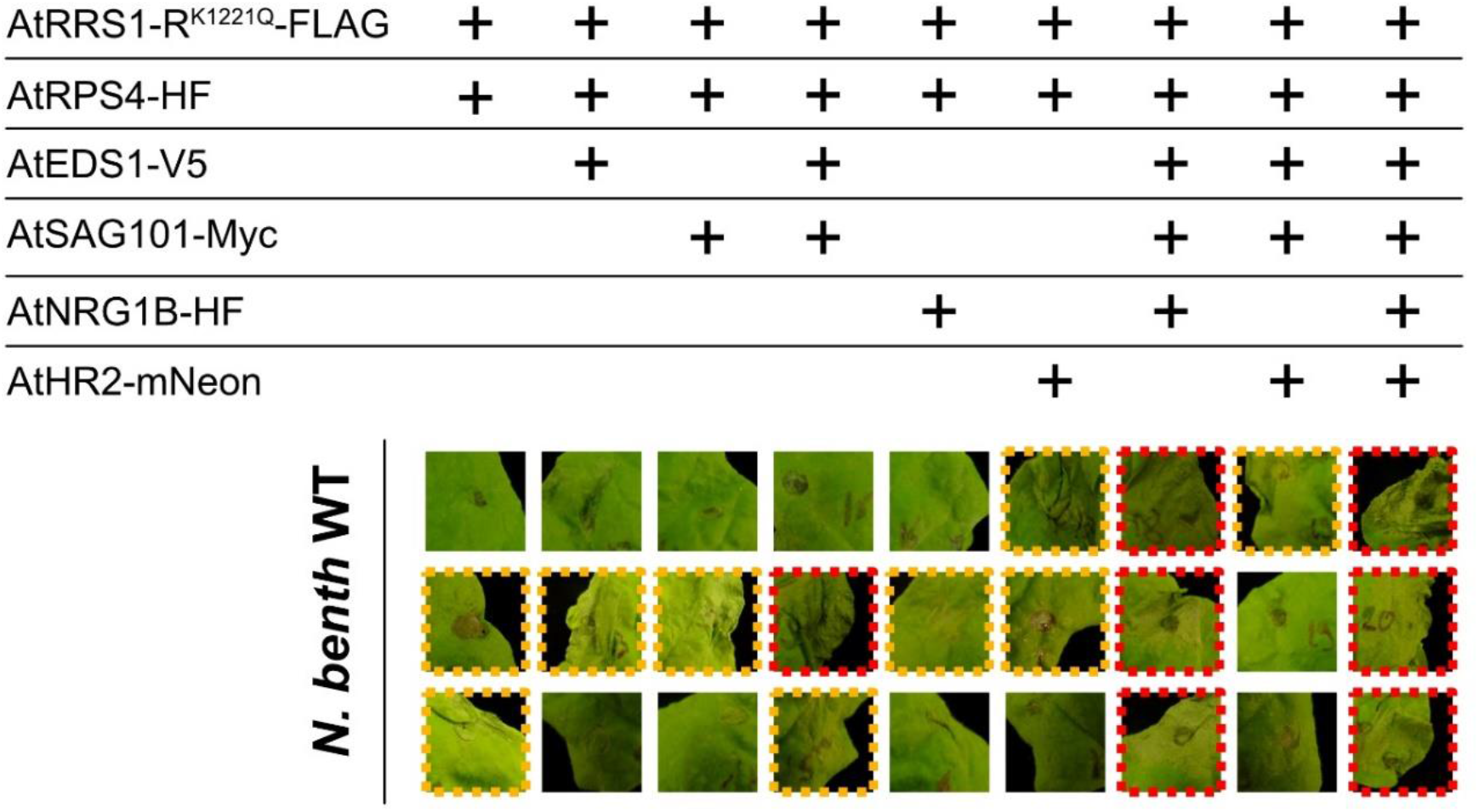
RRS1/RPS4-mediated HR in *N. benthamiana* requires EDS1/SAG101/NRG1, but not HR2. RRS1-R^K1221Q^ (an RPS4-dependent auto-active form of RRS1-R) and RPS4 with EDS1 and/or SAG101 and/or NRG1B and/or HR2 from Arabidopsis, were transiently expressed in *N. benthamiana*. *A. tumefaciens* strains GV3101 carrying denoted construct were infiltrated in 4-week old leaves, OD_600_ = 0.04. Pictures were taken three days after infiltration. Three pictures per combination indicates infiltration of three different plants. Red indicates full HR. Orange indicates partial HR. **Figure S1** displays more control combinations from the same assay.

### Col-0_*rpw8* mutant supports more growth of a virulent bacterial pathogen

RRS1/RPS4-mediated HR is not RPW8-dependent (**Figure 4 and Figure 5**). Consistently, the growth of *P. syringae* pv tomato strain DC3000 carrying AvrRps4 is not affected in Col-0_*rpw8* (**Figure 6A and** **S2A-B**). AvrRps4 avirulence function observed in Col-0, is lost in Col-0_*eds1* but maintained in Col-0_*rpw8*, indicating that RPW8 is not required for RRS1/RPS4-mediated resistance. In contrast, the growth of DC3000 carrying an empty vector is significantly enhanced in Col-0_*rpw8* compared to that in Col-0 (**Figure 6B and S2C-D**). In fact, Col-0_*rpw8* is as susceptible as Col-0_*eds1*, indicating that loss of all RPW8 homologs compromises resistance to bacterial pathogens.

**Figure 6:**
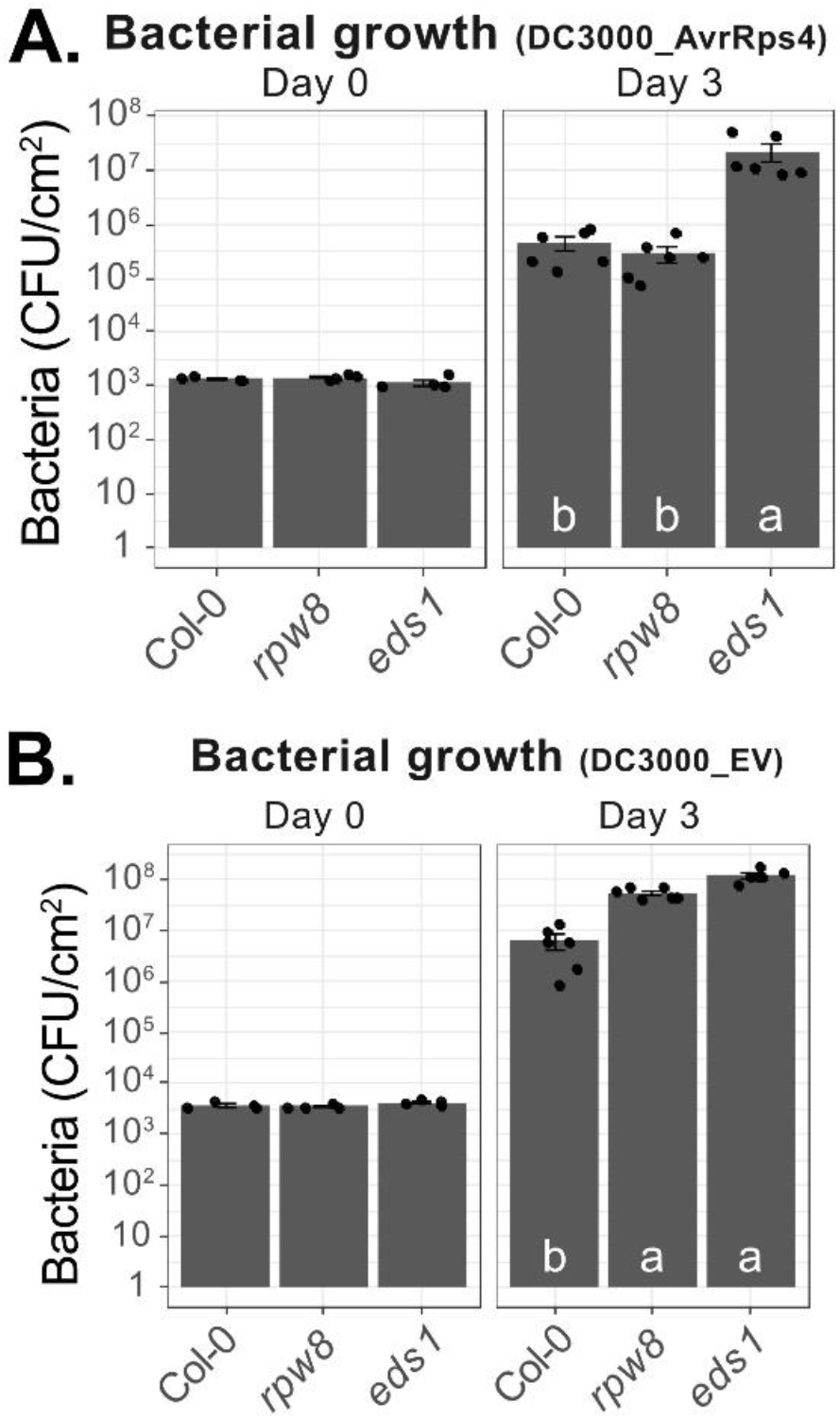
Resistance to an avirulent *P. syringae* strain containing *AvrRps4* is not affected whereas growth of a virulent *P. syringae* strain is enhanced in Col-0_*rpw8*. Five-week old plants were infiltrated with *P. syringae* pv. tomato strain DC3000, carrying AvrRps4 (**A**) or empty vector (EV, **B**) at OD_600_ = 0.0005. Bacterial quantification was performed just after infiltration (Day 0) and at 3 dpi (Day 3). Each dot represents one individual plant. Letters indicate significant differences (P < 0.05) as determined by a one-way ANOVA followed by post hoc Tukey’s honestly significant difference (HSD) analysis. CFU/cm^2^: colony-forming unit per square centimetre of leaf. Two more biological replicates are displayed in **Figure S2**.

### *RPW8* homologs are not required for WRR4A-mediated resistance to *Albugo candida*

*Albugo candida* is an oomycete causing white rust in Brassicaceae. The race Ac2V, isolated from *Brassica juncea* in Canada, can grow on Arabidopsis transgressive segregants or *eds1* mutants (Rimmer *et al.*, 2000; Cevik *et al.*, 2019). Ac2V is resisted by *WRR4A*, *WRR4B* and an unknown recessive or haplo-insufficient *R*-gene in Col-0 (Borhan *et al.*, 2008; Cevik *et al.*, 2019). We tested Col-0, Col-0_*rpw8* and Col-0_*eds1* with *Albugo candida* Ac2V and found that resistance conferred by the aforementioned *R*-genes requires *EDS1* but not *RPW8* (**Figure 7**).

**Figure 7:**
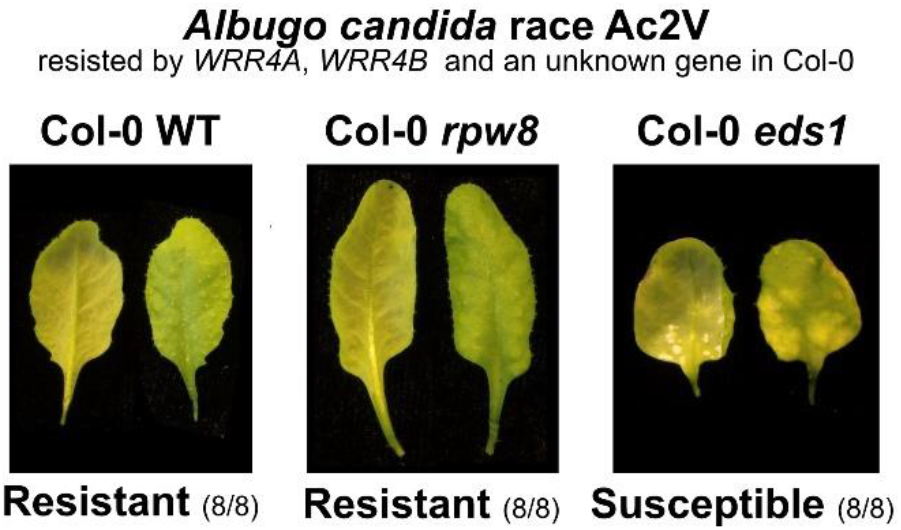
Ac2V resistance does not require *RPW8* in Col-0. 5-week old plants were sprayed inoculated with Ac2V. Plants were phenotyped 12 days after inoculation. Abaxial and adaxial picture of the same leaf are showed. Numbers indicate the number of individual plants showing similar phenotype out of the number of plants tested. Col-0 WT and Col-0_*rpw8* are resistant. Col-0_*eds1* is susceptible.

### Col-0_*rpw8* is more susceptible to adapted and non-adapted powdery mildew fungi

Col-0 is susceptible to the adapted (virulent) powdery mildew isolate *Gc* UCSC1 but still mounts SA-dependent basal resistance against it (Xiao *et al.*, 2005). To test if and how much the four *RPW8* homologs contribute to the basal resistance, we inoculated eight weeks-old plants of Col-0 and Col-0_*rpw8* with *Gc* UCSC1. At 10 dpi, Col-0_*rpw8* plants support more fungal growth than Col-0 plants based on visual scoring in three independent experiments (one representative is shown in **Figure 8A**). Quantification of disease susceptibility showed that Col-0_*rpw8* plants produced ~1.7× spores per mg fresh leaf tissue (**Figure 8B**).

**Figure 8:**
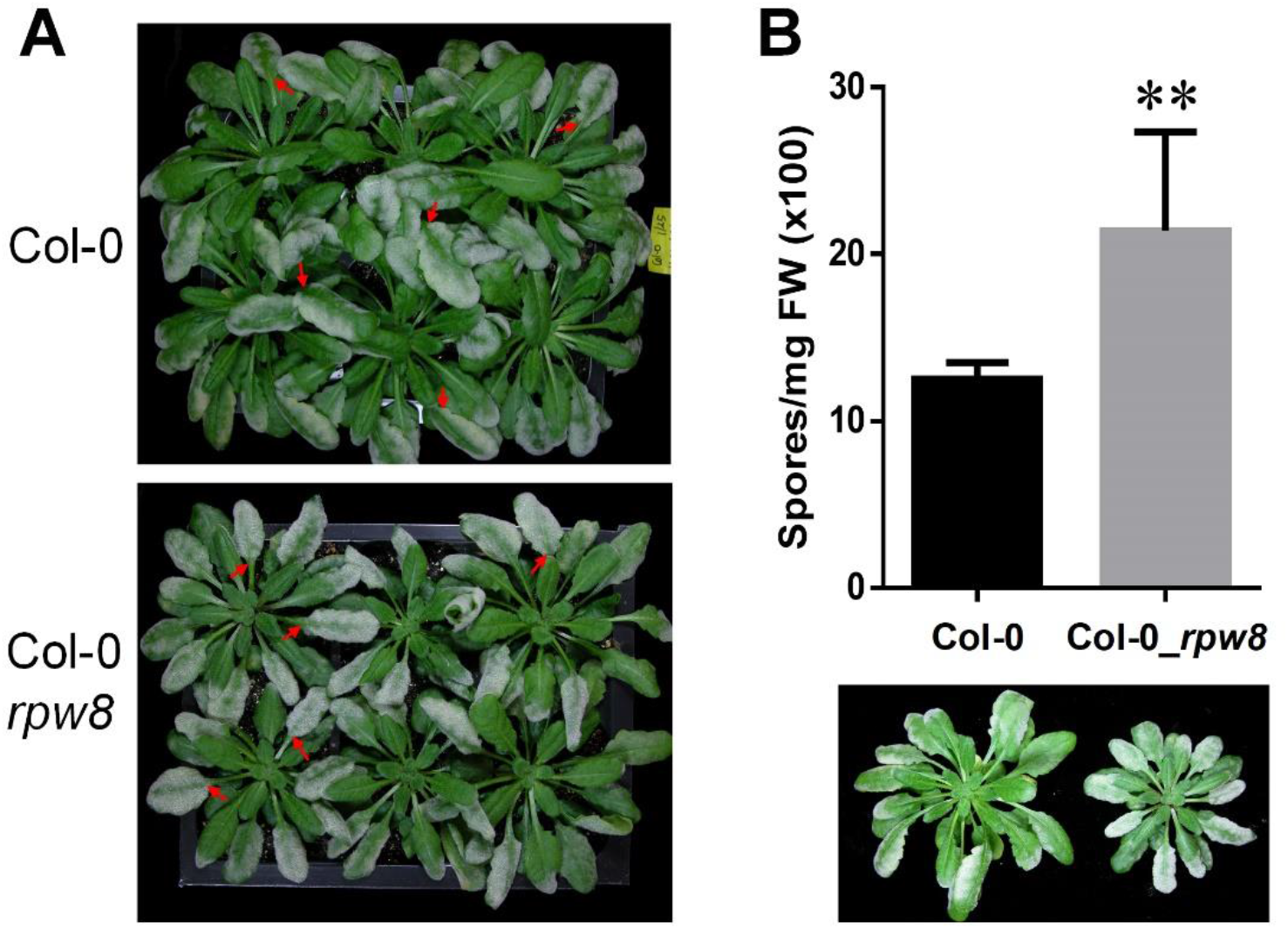
Col-0_*rpw8* is more susceptible to an adapted powdery mildew. **(A)** Eight week-old plants were inoculated with *Gc* UCSC1. Pictures were taken at 10 dpi. Note, leaves of Col-0_*rpw8* were more evenly covered by whitish mildew compared to those of Col-0 (indicated by red arrows). **(B)** Representative infected leaves were weighed and subjected to quantification of total spores per mg fresh leaf tissue. ** indicates significant difference (student *t*-test; P<0.01).

In contrast, Col-0 and 24 other tested Arabidopsis accessions were completely resistant to *Gc* UMSG1, which can only develop very short hyphae and then is arrested shortly after spore germination (Wen *et al.*, 2011). To test if RPW8 homologs are involved in resistance against non-adapted powdery mildew, we inoculated plants of Col-0 and Col-0_*rpw8* with *Gc* UMSG1 and measured total hyphal length at 5 dpi. We found that Col-0_*rpw8* plants supported much more extensive hyphal growth compared to Col-0 (**Figure 9**), indicating that RPW8 homologs in Col-0 collectively contribute to basal resistance against non- or poorly-adapted powdery mildew.

**Figure 9:**
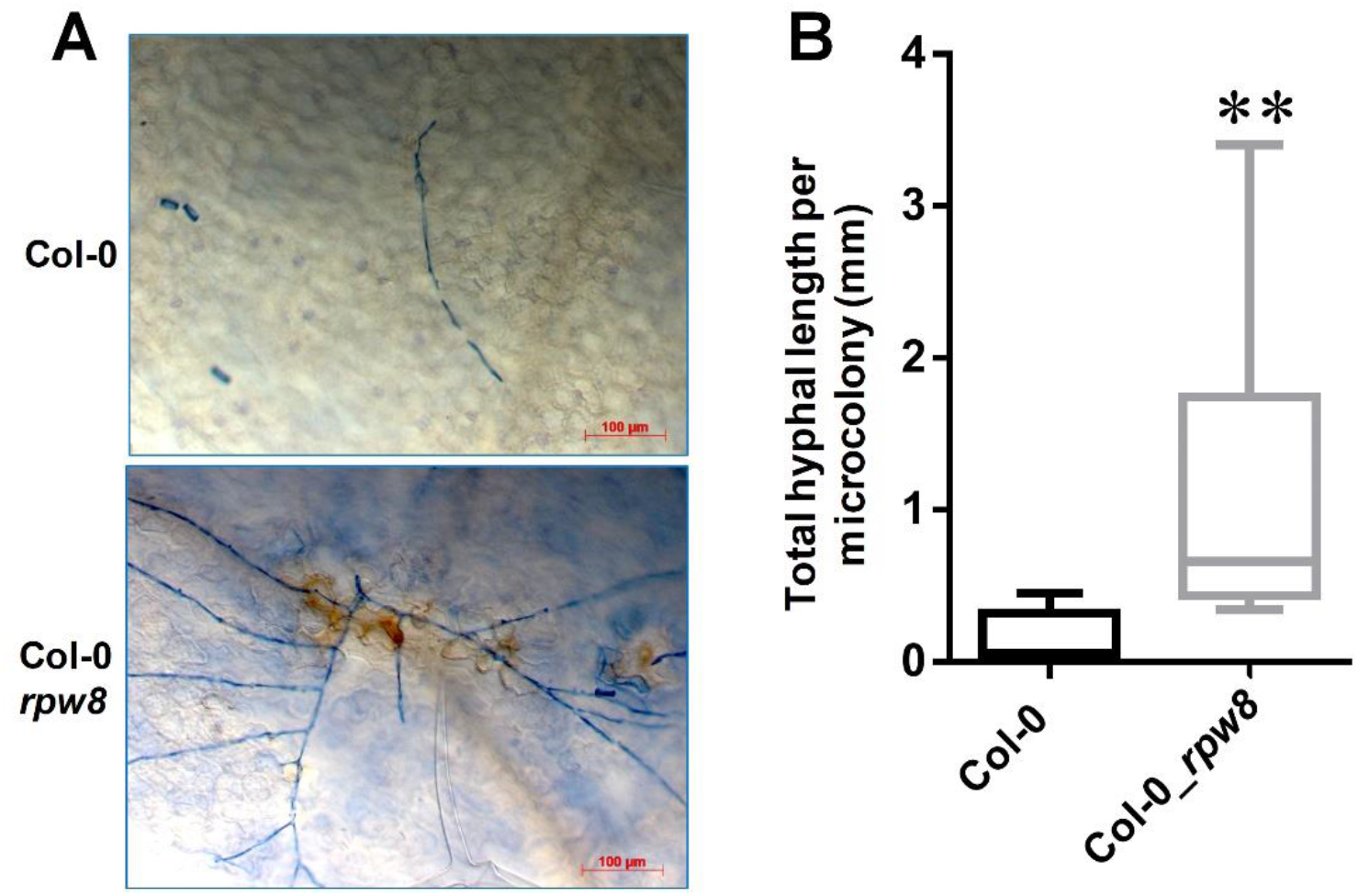
Col-0_*rpw8* supports more hyphal growth of a non-adapted powdery mildew. Eight weeks-old plants were inoculated with *Gc* UMSG1. Leaves were subjected to trypan blue staining at 5 dpi. **(A)** Representative microcolony of the indicated genotypes. **(B)** Total hyphal length of microcolonies. At least 20 microcolonies were measured for total hyphal length for each genotype. ** indicates significant difference (student *t*-test; P<0.01).

## Discussion

Based on previous characterisation of RPW8 (Xiao *et al.*, 2005; Collier *et al.*, 2011; Barragan *et al.*, 2019; Li *et al.*, 2019), we propose the following two hypotheses that could explain the molecular and biochemical function of RPW8. The first hypothesis posits that RPW8 homologs and the RPW8 domain in RPW8-NLRs participate in activation of immune responses through oligomerization and membrane insertion (pore-forming), thus contributing to HR and pathogen resistance. The recent analysis revealing similarities between RPW8 and the HeLo-domain-containing proteins MLKL, HELLP and HET-S from animals and fungus (Daskalov *et al.*, 2016) supports this hypothesis. In animals and fungi, these HeLo-domain-containing proteins can oligomerise (upon sensing of various signals) causing the HeLo-domain to form a disruptive membrane insertion structure (*i.e.* pore-forming), resulting in cell death (Cai *et al.*, 2017). Based on the presence of a putative N-terminal HeLo domain, MLKL-encoding genes have recently been identified in plants (Mahdi *et al.*, 2019). Ablation of all the three *MLKL* genes in Arabidopsis compromised resistance against biotrophic pathogens. These plant MLKL proteins can form tetramers and are associated with microtubules, suggesting a cell death-independent immunity mechanism (Mahdi *et al.*, 2019). How these MLKL proteins enhance plant immunity remains unresolved. It is possible that RPW8 homologs and plant MLKL-like proteins define a HeLo-domain-containing protein family in plants and serve similar functions as their fungal and animal counterparts. If so, RPW8-NLRs such as ADR1 and NRG1 could then be renamed HeLo-NLRs. The earlier observations that RPW8.2 and other RPW8 homologs are localized to the EHM (Wang *et al.*, 2009; Berkey *et al.*, 2017) and over-expression of RPW8 results in massive cell death (Xiao *et al.*, 2003) support this hypothesis. However, the precise molecular function of the putative N-terminal HeLo-domain of MLKL and RPW8 has yet to be characterized.

In this study, we showed that the HR triggered by five well-described NLRs does not require any RPW8 homologs in Arabidopsis **(Figure 4)**. In addition, HR2 is not sufficient to reconstruct RRS1/RPS4-triggered HR in *N. benthamiana* (**Figure 5**). By contrast, loss of all four *RPW8* homologs compromised resistance against *P. syringae* and powdery mildew (**Figure 6**, **Figure 8 and Figure 9**). These observations, together with the earlier findings that RPW8 enhances resistance against an oomycete (*Hpa)* and interacts with RPP7 (Wang *et al.*, 2007; Li *et al.*, 2019), suggest that RPW8 plays an important role in plant immunity but is dispensable for the HR. However, unlike plant MLKLs whose role in cell death is unclear (Mahdi *et al.*, 2019), overexpression of wild-type RPW8 homologs or variants or C-terminal YFP-tagged HR3 results in cell death (Xiao *et al.*, 2003; Wang *et al.*, 2013; Berkey *et al.*, 2017). Since all tested RPW8 homologs are capable of membrane targeting during infection (Wang *et al.*, 2013; Berkey *et al.*, 2017), RPW8 function could be to direct the associated protein complex to a specific subcellular compartment to activate local immune response, including cell death in extreme cases.

According to the second hypothesis, RPW8 proteins are decoys for RPW8-NLR-targeting effectors. However, such avirulent effectors are not known. Even if they exist, the loss of their function in Col-0_*rpw8* would not been seen if other avirulent effectors get recognised by an RPW8-independent mechanism. Given that the pathogens tested in our study contain up to several hundreds of effectors, this may well be the case. Thus, our results neither validate or invalidate this RPW8 decoy hypothesis. Characterisation of an RPW8-targeted effector would enable further investigation of this question.

A detailed search for RPW8 and RPW8 domain revealed 11 predicted proteins in Col-0 (Zhong & Cheng, 2016). Four are Homologues of RPW8 (HR1, HR2, HR3 and HR4), five are RPW8-containing NLRs (ADR1, ADR1-L1, ADR1-L2, NRG1A and NRG1B), one is a non-canonical NLR (DAR5: RPW8-NB-ARC-LIM) and the last one is encoded by *AT3G26470*. The predicted protein encoded by *AT3G26470* has 221 amino acid protein (25.3 kDa) and contains an N-terminal RPW8 domain. *AT3G26470* displays a two-exons structure, typical for *RPW8*. We thus consider this gene a distant Homologue of RPW8 and called it HR5. However, the RPW8 domain of HR5 resembles more the RPW8 domain of ADR1-L1 than that of NRG1 or HR1, HR2, HR3 and HR4 (**Figure S3**). Hence, HR5 might have resulted from a recent duplication event of the ADR1-L1 RPW8 domain. Similar to the above, HR5 could be a decoy for an ADR1-L1-targeting effector, independently of any HR1, HR2, HR3 and/or HR4 function. Parsimony suggests that HR1, HR2, HR3 and HR4 share a common ancestor, which is not shared with HR5. Thus, if HR1, HR2, HR3 and HR4 play a redundant function, it is not likely to be shared with HR5.

A recent study highlighted co-evolution between EDS1, SAG101 and NRG1 within plant clades to regulate downstream pathways in TIR-NLR-mediated immunity. Particularly, these three components need to have co-evolved to be functional (Lapin *et al.*, 2019). In this study, we tested whether HR2 is part of the minimal required components to reconstruct RRS1/RPS4-triggered HR in *N. benthamiana*. We found that co-expression of Arabidopsis EDS1, SAG101 and NRG1 is necessary to reconstitute RRS1/RPS4-triggered HR in *N. benthamiana*, but HR2 is dispensable (Fig. 5). Thus, RPW8 is neither required nor sufficient, so likely not involved, in RRS1-RPS4-mediated HR. We confirmed that EDS1, SAG101 and NRG1 need to come from the same genome to function, as previously reported (Lapin *et al.*, 2019). However, unlike Roq1 (a TIR-NLR from *N. benthamiana*) that can signal via EDS1/SAG101/NRG1 from either Arabidopsis or *N. benthamiana* genomes (Lapin *et al.*, 2019), RRS1-RPS4 can only fully signal via Arabidopsis EDS1/SAG101/NRG1. It indicates that the TIR-NLRs Roq1 and RPS4 are sensed differentially by EDS1, SAG101 and NRG1. Recently, TIR domains of plant NLRs have been shown to possess enzymatic activity (Horsefield *et al.*, 2019; Wan *et al.*, 2019). They can degrade nicotinamide adenine dinucleotide in its oxidized form (NAD+) into adenine dinucleotide ribose (ADPR), variant-cyclic ADPR (v-cADPR) and nicotinamide (Nam) (Wan *et al.*, 2019). Our data show that *N. benthamiana* EDS1/SAG101/NRG1 does not sense RRS1/RPS4 activity, but Arabidopsis EDS1/SAG101/NRG1 can, even in the *N. benthamiana* system. This suggests a more complex signal transduction pathway between the activation of TIR-NLRs, enzymatic activity and activation of EDS1/SAG101/NRG1.

In conclusion, we generated an *hr1-hr2-hr3-hr4* quadruple mutant (called *rpw8* mutant) in Arabidopsis Col-0. NLR-mediated phenotypes tested remain intact in the mutant. However, Col-0_*rpw8* is partially compromised in resistance against powdery mildew and *P. syringae*, and surprisingly in flowering under short-day growth conditions. Future research is needed to characterize the precise mechanism by which RPW8 proteins or domains contribute to immunity, in comparison to the HeLo-containing proteins in fungi and animals.

## Supporting information

Plasmids

## Acknowledgments

Work at The Sainsbury Laboratory (BC and JDGJ) was supported by the Gatsby Foundation (http://www.gatsby.org.uk/). Work at the University of Maryland (YW and SX) was supported by a National Science Foundation grant (IOS-1457033). The authors thanks Mark Youles (Synbio at The Sainsbury Laboratory) for help in plasmid vector development.

## Supplemental information

**Figure S1:**
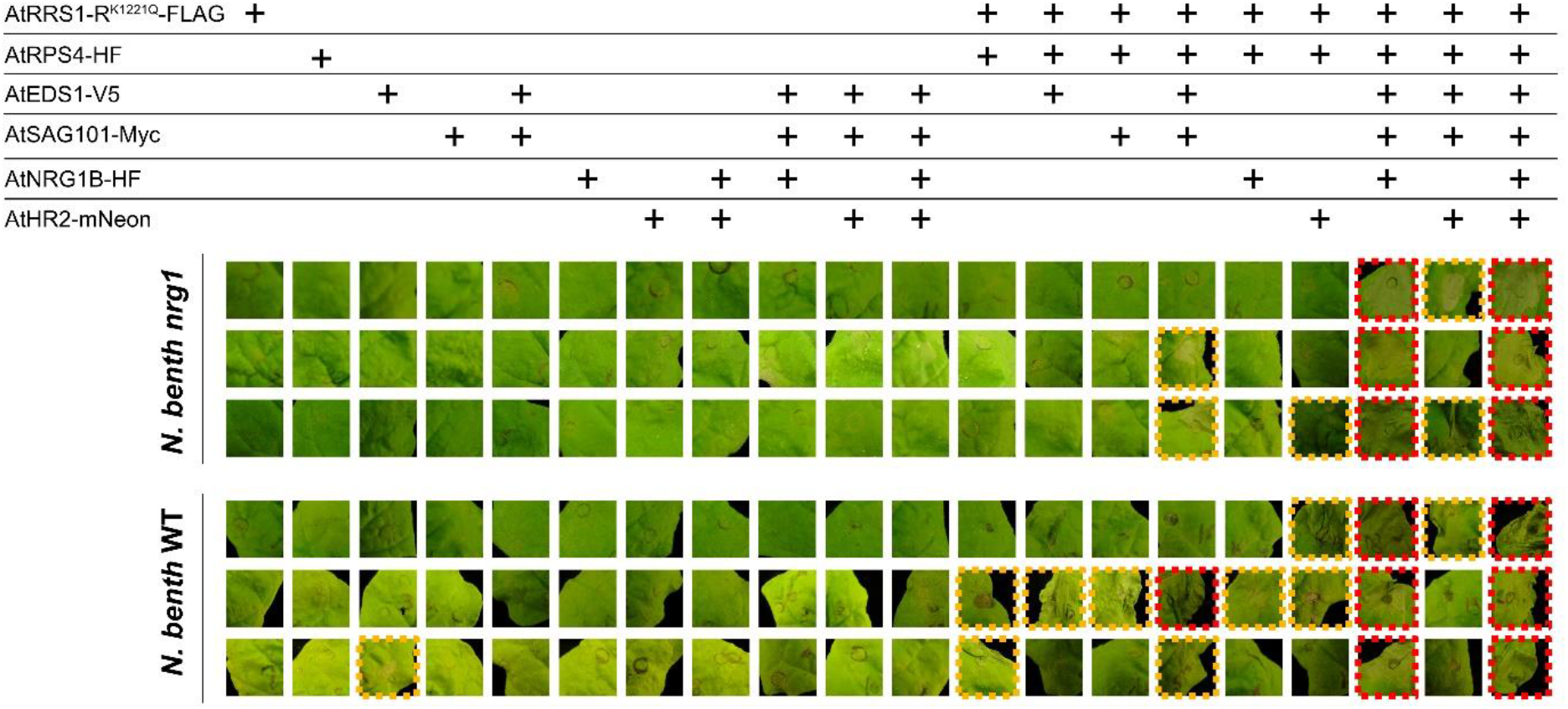
RRS1/RPS4-mediated HR in *N. benthamiana* requires EDS1/SAG101/NRG1, but not HR2. RRS1-R^K1221Q^ (an RPS4-dependent auto-active form of RRS1-R) and RPS4 with EDS1 and/or SAG101 and/or NRG1B and/or HR2 from Arabidopsis, were transiently expressed in 4-week old *N. benthamiana*. *A. tumefaciens* strain GV3101 carrying denoted construct were infiltrated in 4-week old leaves, OD_600_ = 0.04. Pictures were taken 3 dpi. Three pictures per combination indicates infiltration of three different plants. Red indicates full HR. Orange indicates partial HR. **Figure 5** is a partial summary of this assay.

**Figure S2:**
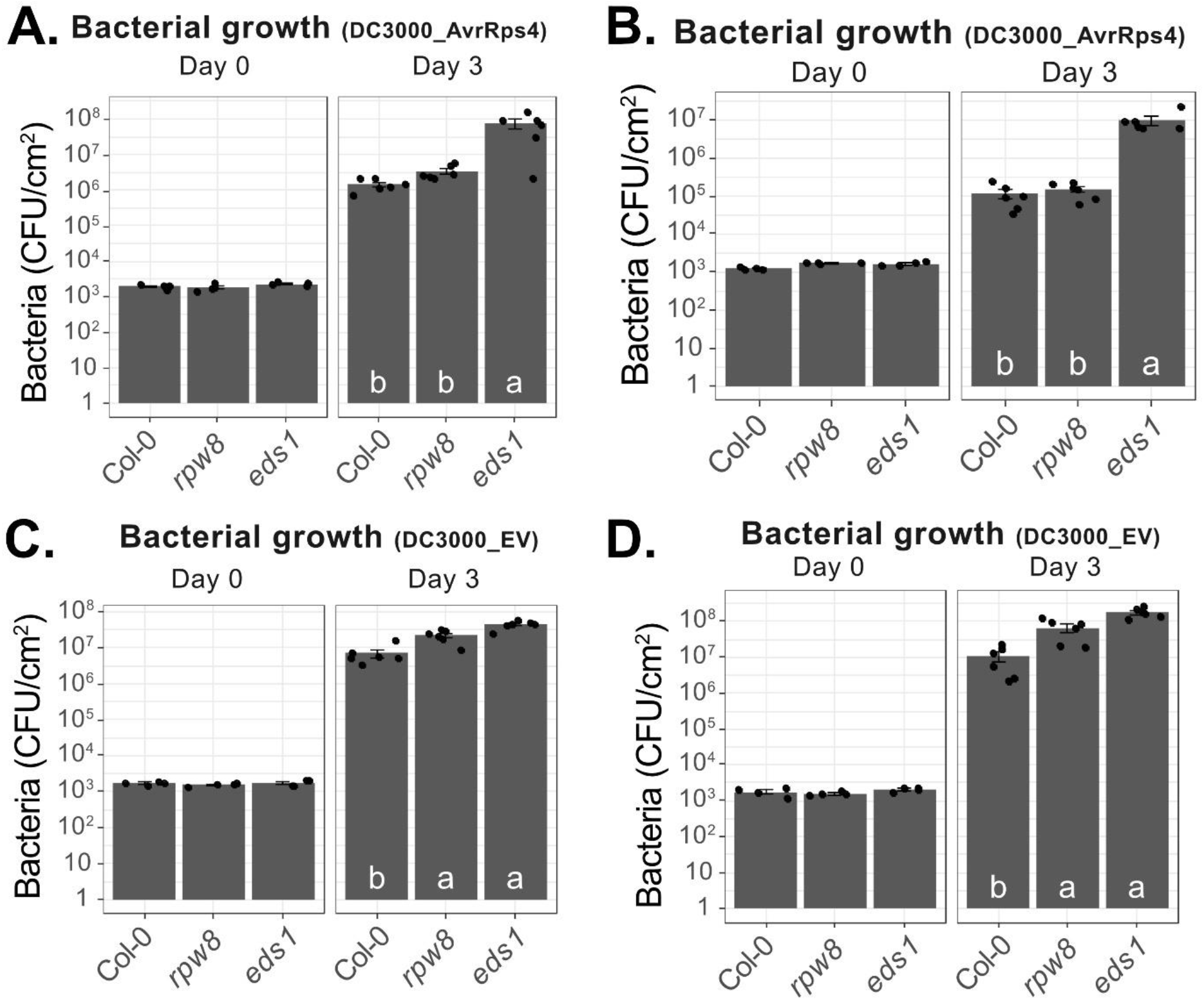
*P. syringae* growth is RPW8-dependent, unlike AvrRps4 avirulence function. Five-week old plants were infiltrated with *P. syringae* pv. tomato strain DC3000, carrying AvrRps4 (**A. and B.**) or empty vector (EV, **C. and D.**) at OD_600_ = 0.0005. Bacterial quantification was performed just after infiltration (Day 0) and at 3 dpi (Day 3). Each dot represents one individual plant. Each panel represent an independent experiment (plant grown at different time). Letters indicate significant differences (P < 0.05) as determined by a one-way ANOVA followed by post hoc Tukey’s honestly significant difference (HSD) analysis. CFU/cm^2^: colony-forming unit per square centimetre of leaf. This figure displays biological replicates of **Figure 6**.

**Figure S3:**
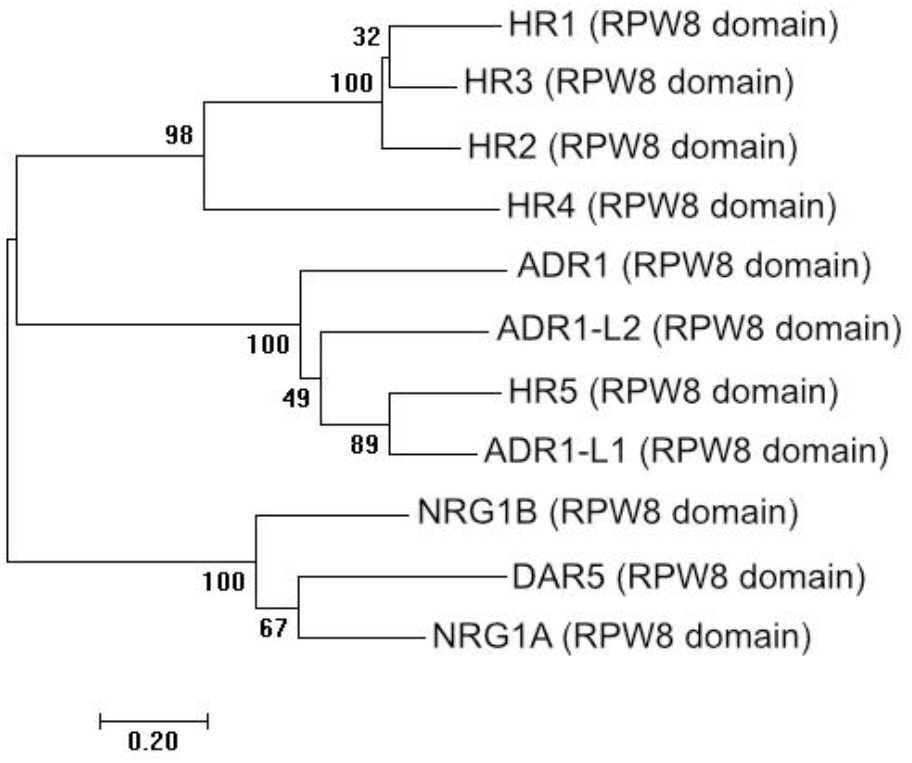
A fifth RPW8 homolog is related to ADR1-L1 RPW8 domain. Neighbor-joining phylogeny of the amino acid sequences of RPW8 domains (identified with SMART, http://smart.embl-heidelberg.de/) from all RPW8-containing proteins in Col-0. Number indicates value of bootstrap 100. The tree is drawn to scale, with branch lengths in the same units as those of the evolutionary distances used to infer the phylogenetic tree. Evolutionary distances are expressed in number of amino acid substitutions per site (Poisson correction method). Tree made using the software MEGA7.

